# Characterization of DNA methylation in PBMCs and donor-matched iPSCs shows methylation is reset during stem cell reprogramming

**DOI:** 10.1101/2024.12.13.627515

**Authors:** Xylena Reed, Cory A. Weller, Sara Saez-Atienzar, Alexandra Beilina, Sultana Solaiman, Makayla Portley, Mary Kaileh, Roshni Roy, Jinhui Ding, A. Zenobia Moore, D. Thad Whitaker, Bryan J. Traynor, J. Raphael Gibbs, Sonja W. Scholz, Mark R. Cookson

## Abstract

Graphical Abstract

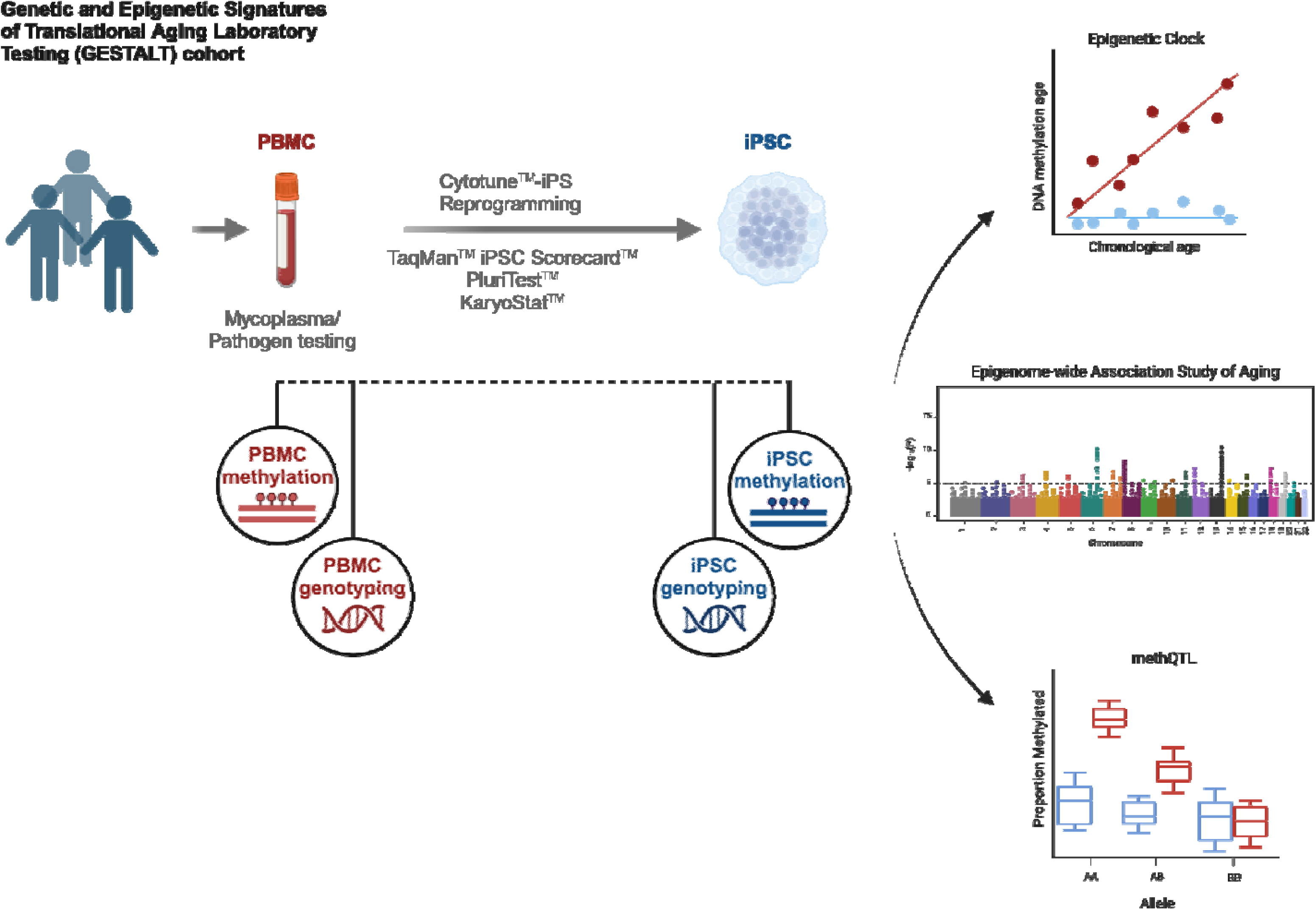

**Highlights:**

- Generation of a population-level set of iPSC lines from healthy individuals across the lifespan
- Aging-related features were reset based on epigenetic markers of cytosine methylation and telomere length
- By comparing methQTLs in iPSC vs. their donor PBMCs, we find that detection of methQTLs reflect biological functions of different cell types

DNA methylation is an important epigenetic mechanism that helps define and maintain cellular functions. It is influenced by many factors, including environmental exposures, genotype, cell type, sex, and aging. Since age is the primary risk factor for developing neurodegenerative diseases, it is important to determine if aging-related DNA methylation is retained when cells are reprogrammed to an induced Pluripotent Stem Cell (iPSC) state. Here, we selected peripheral blood mononuclear cells (PBMCs; n = 99) from a cohort of diverse and healthy individuals enrolled in the Genetic and Epigenetic Signatures of Translational Aging Laboratory Testing (GESTALT) study to convert to iPSCs. After reprogramming we evaluated the resulting iPSCs for DNA methylation signatures to determine if they reflect the confounding factors of age and environmental factors. We used genome-wide DNA methylation arrays in both cell types to show that the epigenetic clock is largely reset to an early methylation age after conversion of PBMCs to iPSCs. We further examined the epigenetic age of each cell type using an Epigenome-wide Association Study (EWAS). Finally, we identified a set of methylation Quantitative Trait Loci (methQTL) in each cell type. Our results show that age-related DNA methylation is largely reset in iPSCs, and each cell type has a unique set of methylation sites that are genetically influenced.

## Introduction

Methylation of cytosine residues is a stable epigenetic modification of DNA seen across species and is responsible for controlling many aspects of gene expression (Law and Jacobsen, 2010). In mammalian cells, cytosine is methylated predominantly at CpG sites by the enzyme DNA methyltransferase 1. While many CpG sites in the genome show high levels of constitutive methylation, the proportion of DNA methylated at other sites depends on the cellular differentiation state and other environmental and biological effects, including aging (D’Aquila et al., 2013; Ferrucci et al., 2020; Marioni et al., 2019). DNA methylation also correlates with genetic variation, leading to the establishment of methylation quantitative trait loci (methQTL), similar to expression quantitative trait loci (eQTL) for RNA expression (Gibbs et al., 2010; Nabais et al., 2023; Villicaña and Bell, 2021; Zhang et al., 2010). Thus, the overall proportion of CpG methylation in each tissue reflects a complex interplay between genetic and non-genetic influences.

Induced pluripotent stem cells (iPSC) are widely used to model cell biological events without cell transformation. For example, iPSC models have been used to replicate tissue-based methQTL and eQTL in an endogenous genomic context, suggesting that these models retain genetically encoded influences on epigenetics and gene expression (Cuomo et al., 2020; Salles et al., 2024). However, there has been some disagreement on whether iPSC models retain epigenetic marks that arise from cellular differentiation or organismal aging, as these are rewritten during reprogramming (Horvath, 2013; Lo Sardo et al., 2017; Miller et al., 2013). While some types of DNA methylation may be lost, iPSCs can be grown under consistent conditions and so, by inference, the contribution of environmental factors should be consistent between different donor lines. Thus, iPSC models may be useful in identifying methQTL independent of other sources of methylation variation (Ruiz et al., 2012).

Here, we generated a new collection of iPSC lines from a cohort of healthy donors, ranging from 22 to 92 years old, to examine the effects of cellular reprogramming on CpG methylation. We first obtained peripheral blood mononuclear cells (PBMCs) collected by apheresis from 99 individuals in the ongoing Genetic and Epigenetic Signatures of Translational Aging (GESTALT) longitudinal study that aims to identify biomarkers related to aging (Roy et al., 2023, 2021; Tsitsipatis et al., 2023, 2022; Tumasian et al., 2021). The strict inclusion criteria for the study’s participants result in the retention of healthy individuals, allowing us to examine CpG methylation signatures in the absence of disease. Additionally, the availability of genotyping and methylation data from the same donors before and after reprogramming allowed us to compare methQTL between matched iPSCs and PBMCs.

We show that iPSCs lack age-related methylation signatures tested using five different epigenetic clocks and an Epigenome-wide Association Study (EWAS). Additionally, CpG methylation patterns are highly distinct between iPSCs and PBMCs, and while we recovered abundant evidence of genetic effects on CpG methylation in both PBMCs and iPSCs, some of which were correlated between cell types, we found that there is a unique set of methQTL for each cell type. We found that methQTL in PBMCs are enriched for gene associations that are involved in biological processes involving lipid localization and steroid secretion.

## Results

### Generation of a population resource for iPSC from diverse, healthy donors across the age span

Our overall strategy to address the relative contributions of aging, environment, genetics and cell type on DNA methylation is shown in Fig. 1A. We first set out to generate a series of iPSC lines from PBMCs collected from 99 healthy individuals in the GESTALT cohort (Table 1 and Supplementary Table 1). After testing PBMCs for mycoplasma and human pathogens, cells were reprogrammed using Cytotune delivered by Sendai virus, with two clones retained for each donor. Of the starting 99 initial donors, 93 lines were effectively reprogrammed (Supplementary Fig. 1A). Quality of reprogramming was evaluated using a TaqMan-based iPSC Scorecard (Bock et al., 2011), PluriTest, and KaryoStat (Supplementary Fig. 1B and Supplementary Table 2), and all lines passed conventional thresholds for each test.

**Figure 1.**
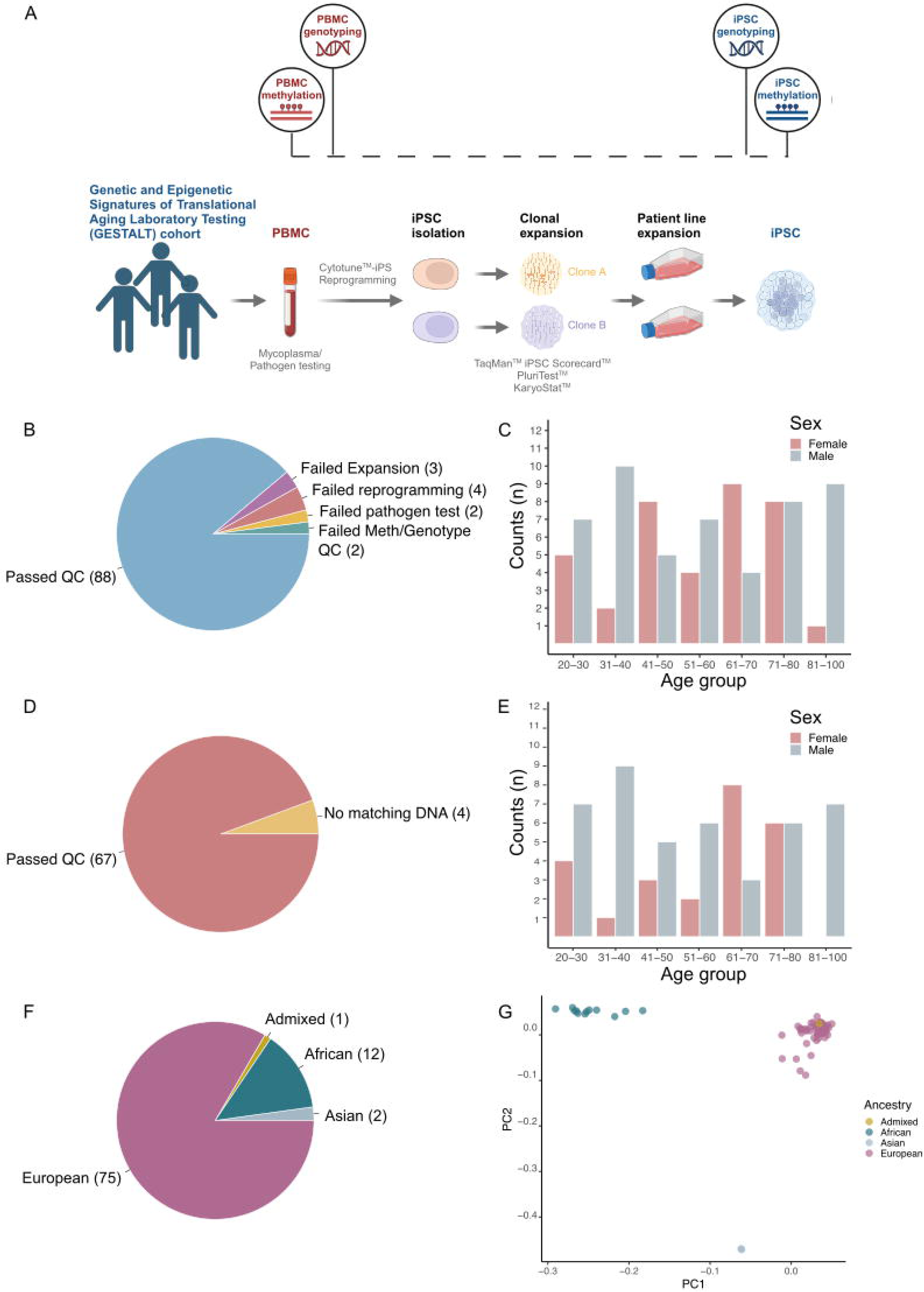
Experimental workflow and sample characteristics in a healthy donor series. **A**. Experimental workflow. **B**. Number of iPSC lines passing each data quality control threshold, and showing those that failed reprogramming, expansion, and methylation or genotype quality control checks. **C.** iPSC lines that passed QC binned by age and sex. **D**. Number of PBMC donor methylation samples passing data quality thresholds, and showing those without donor matched iPSC DNA or failed DNA methylation QC checks. **E.** PBMC samples that passed QC checks binned by age and sex. **F**. Number of samples remaining by ancestry group after QC. **G.** Genetic principal components analysis (PCA) of remaining samples compared to the HapMap 3 Genome Reference Panel.

**Table 1.** Demographic data of the iPSC and PBMC lines that passed Quality Control (QC). Sample size indicates number of individuals in each category, and parenthesis indicates the percentage of individuals in each category.

|  |  | iPSC | PBMC (%) |
| --- | --- | --- | --- |
| **Gender, n (%)** | **Female** | 39 (43.33) | 24 (35.8) |
|  | **Male** | 51 (56.67) | 43 (64.2) |
| **Age at collection, mean (SD)** |  | 55.17 (19.47) | 54.37 (20.03) |
| **Ancestry, n (%)** | **Admixed** | 1(1.11%) | 1 |
|  | **African** | 12 (13.33%) | 10 (10.4) |
|  | **Asian** | 2 (2.22%) | 2 (3.0) |
|  | **European** | 75 (83.3%) | 57 (74.6) |
| **Total (n)** |  | 90 | 67 |

Of the 93 reprogrammed cell lines, 90 were successfully expanded. Following clone expansion, each line was genotyped on an Infinium Global Diversity Array (Illumina). We confirmed the identity of each line by comparing iPSC genotypes to genotypes previously generated from the PBMCs of each donor, and two lines were discarded for failing methylation and genotyping.

After quality control analysis, the iPSC cohort consisted of 88 total lines made up of individuals of European (n = 75), Black or African American (n = 12), and Asian ancestries (n = 2), as well as one individual who self-reported multiple ethnicities (based on genetic PCs this individual maps with the European samples in Fig. 1G) (Fig. 1 F, G; Table 1, Supplementary Table 1).

Thus, we successfully generated a cohort of 88 iPSC lines from healthy donors containing individuals with differing genetic ancestry.

### Cell state and inter-individual differences contribute to methylation status

We generated genome-wide methylation data from both clones for each iPSC line using the Illumina Infinium MethylationEPIC v2.0 array and performed stepwise quality control checks at sample- and probe-level after normalization (Materials and Methods). Methylation data from the EPIC array were also available from a subset of the original PBMC samples (n = 73). CpG methylation values showed a bimodal distribution for both PBMCs and iPSCs, as expected.

After quality control, we retained 67 donor-matched iPSC-PBMC pairs and compared the resulting methylation signatures from the same donors across iPSC clones and between cell types (Fig. 1D, E; Table 1). We calculated the Pearson correlation coefficient between samples using methylation probes that were informative for both cell types, i.e. probes within the top 5000 most variable probes for both iPSCs and PBMCs. We found that the iPSC replicate clone-to-clone correlation (ρ=0.911 ± 0.057, number of samples with two clones = 70) was higher than ρ the PBMC-to-iPSC correlation within the same donor (ρ =0.714 ± 0.039, Table 2, Supplementary Fig. 2). Both measures were significantly higher than the mean correlation between iPSCs from different donors (ρ=0.522 ± 0.0343). All group comparisons were significantly different from each other (ANOVA: *df*=4, *F*=10320, *p*<2×10^−16^; Tukey HSD, all comparisons: p-value < 2×10^−16^). We also performed a correlation analysis of DNA methylation against all PBMCs-iPSC clones for cells from different donors and saw a much lower correlation (ρ=0.383 ± 0.055; Supplementary Fig. 3). Collectively, these data show that inter-individual differences contribute to methylation status, but also that a significant source of variability for DNA CpG methylation relates to cell state.

**Table 2.** Methylation correlation between and within donors and cell types.

| **Comparison** | **Cells** | **Number of comparisons** | **Mean** | **SD** |
| --- | --- | --- | --- | --- |
| Same Donor | iPSC to iPSC | 70 | 0.911 | 0.056 |
| Same Donor | iPSC to PBMC | 128 | 0.714 | 0.039 |
| Different Donor | iPSC to iPSC | 12491 | 0.522 | 0.073 |
| Different Donor | PBMC to PBMC | 2485 | 0.578 | 0.050 |
| Different Donor | iPSC to PBMC | 11161 | 0.383 | 0.055 |

### iPSC clones reset known epigenetic markers of aging

We next evaluated whether iPSC lines are epigenetically young relative to their donors in this healthy population study across the lifespan. We estimated the epigenetic age using the Horvath multi-tissue predictor of epigenetic age, which considered the methylation state of 353 CpGs (Horvath, 2013). As expected, PBMC DNA methylation was strongly correlated with donor age (r = 0.905, p-value = 2.95 x 10^−37^, n = 67; Fig. 2A). The estimated average methylation age was 53.33 ± 16.75 years. In contrast, iPSC DNA methylation showed no correlation with donor age (r = 0.09, p = 2.24 x 10^−3^, n = 70), and the estimated methylation average age was −0.24 ±0.17 years. These results were confirmed with additional DNA methylation clocks, including the clock developed by Hannum et al. (Fig. 2B), the phenoAge clock (Fig. 2C), and two age predictors described by Zhang et al. based on best linear unbiased prediction (BLUP; Fig. 3D), and elastic net prediction (EN; Fig. 2E) (Hannum et al., 2013; Levine et al., 2018; Zhang et al., 2019). In all analyses, the epigenetic clocks showed no correlation between methylation age and chronological age in iPSCs, while the corresponding donor PBMCs displayed a strong correlation, indicating that the loss of epigenetic aging markers during reprogramming was robust across different measures (Supplementary Table 3).

**Figure 2:**
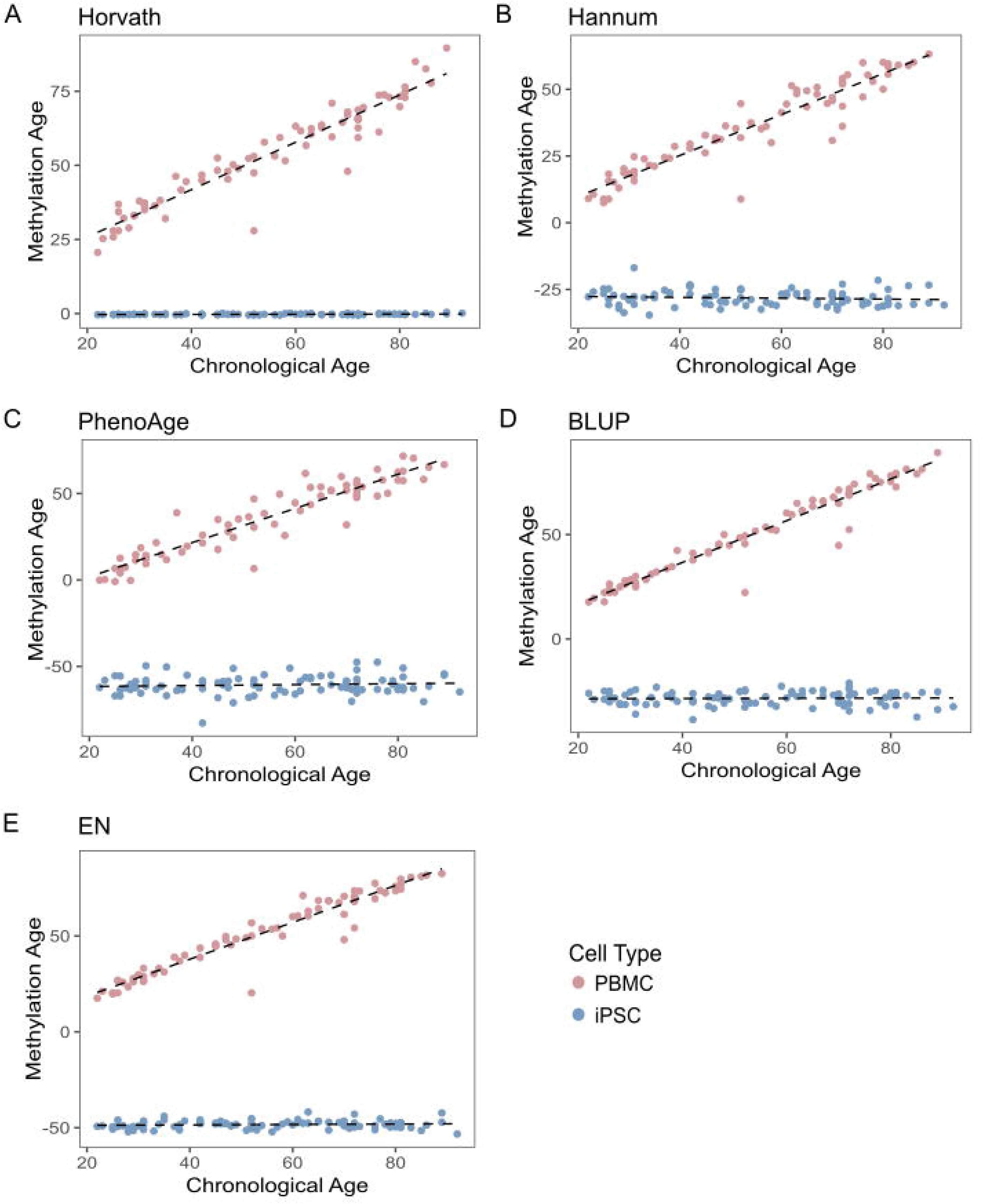
Comparison of methylation aging signatures in PBMCs (red) and iPSCs (blue). Plots showing chronological (x-axis) versus methylation age (y-axis) calculated using different epigenetic clocks. Each point corresponds to one donor sample. The red points represent DNA methylation from PBMCs, and the blue points show DNA methylation from iPSCs. **A**. Using the Horvath epigenetic clock, the correlation values between chronological and methylation age in PBMCs is 0.905 (p-value = 2.95×10^−37^) while the correlation for iPSCs is 0.09 (p-value = 0.00224). **B**. Using the Hannum clock, the correlation values between chronological and methylation age in PBMCs is 0.884 (p-value = 3.62×10^−34^) and iPSCs is (p-value = 0.312) **C**. Using the PhenoAge clock, the correlation values between chronological and methylation age in PBMCs is 0.875 (p-value = 4.70×10^−33^) and the iPSC correlation is −0.002 (p-value = 0.368) **D**. Using the BLUP clock, the correlation values between chronological and methylation age in PBMCs is 0.936 (p-value = 4.78×10^−43)^ and the correlation in iPSCs is −0.01 (p-value = 0.789) **E**. Using the EN clock, the correlation values between chronological and methylation age in PBMCs is 0.923 (p-value = 2.38×10^−40^) and in iPSCs is 0.001 (p-value = 0.299).

**Figure 3.**
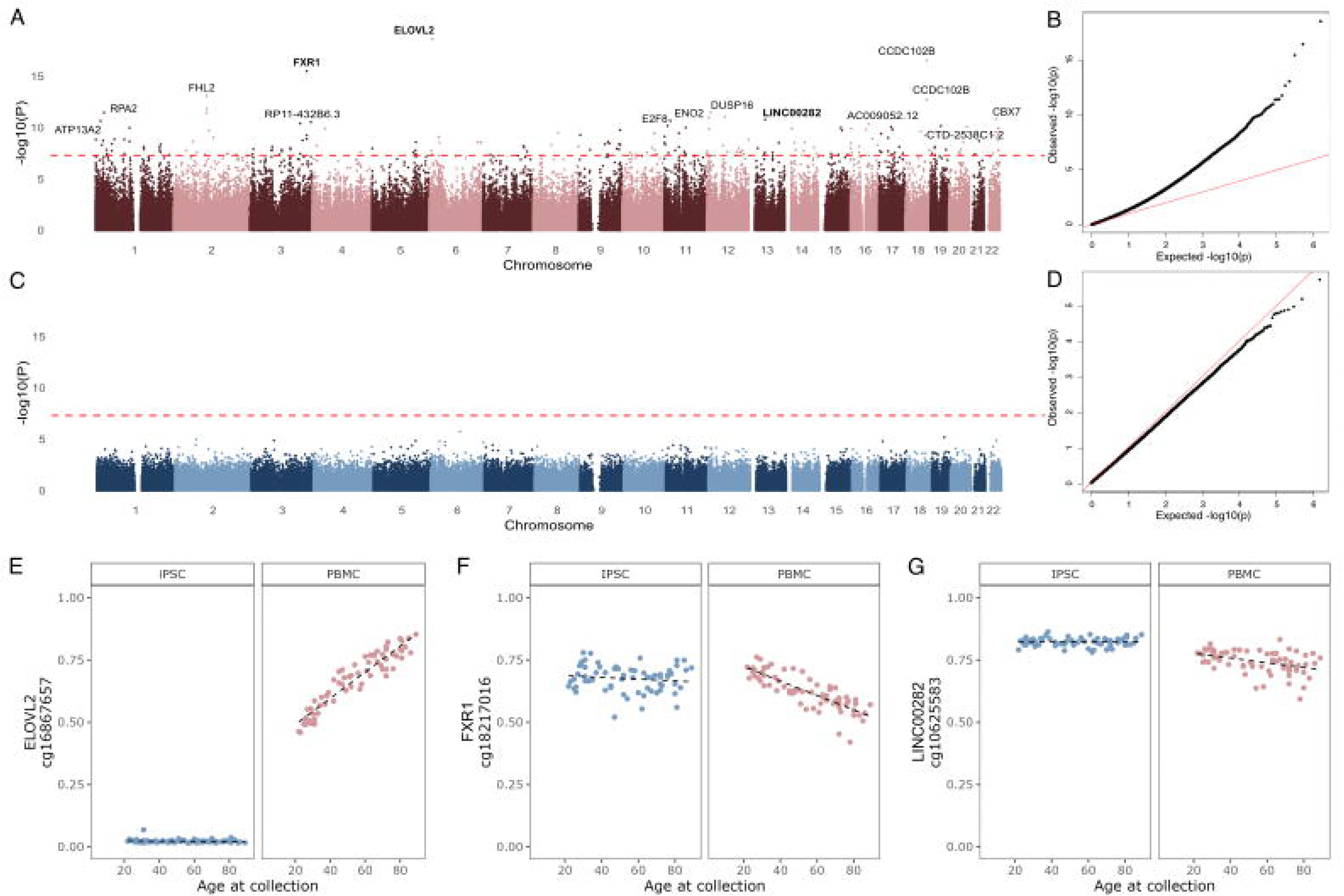
Epigenome-wide Association Study (EWAS) in PBMCs and reprogrammed iPSCs. EWAS identified CpGs associated with the donor’s age at the time of collection in PBMCs (red) and iPSCs (blue). **A**. Manhattan plot showing significant associations between DNA methylation at common CpG sites and age in PBMCs. **B.** The association analysis quantile-quantile (QQ) plots corresponding to PBMCs. The expected p-values are on the x-axis, and the observed p-values are on the y-axis. P-values are transformed to a -log10 scale. **C**.The Manhattan plots shows the lack of significant associations (dashed red line = p-value < 5 x 10^−8^) between DNA methylation levels of common CpG sites and age in iPSCs. **D.** QQ plot corresponding to the iPSC EWAS. **E-G.** Representative examples of significant DNA methylation CpGs associated with donor age at the time of collection based on EWAS results in PBMCs. The individual points represent the methylation level per individual measured in PBMCs (red) or iPSCs (blue). Probes were mapped to gene names using the EPICv2 hg38 annotation manifest.

DNA methylation clocks are based on multiple regression models across various informative genomic sites. As a result, we hypothesized there might be residual aging signals outside of the tested groups of CpG methylation sites. We therefore considered whether other individual DNA CpG methylation sites may show an association with donor age and performed an Epigenome-Wide Association Study (EWAS) using donor age as an outcome measure in PBMCs and iPSCs. We identified aging-associated changes in methylation in PBMCs, yet recovered no significant associations within the iPSC data (Fig. 3A-D). For PBMCs, while we replicated some sites already associated with aging by one or more methylation clocks (for example the most significant site overall, cg1687657 near ELOVL2 (Fig. 3E)), we also identified new sites not previously associated with aging. For example, two newly identified sites, cg18217016 (FXR1) and cg10625583 (LINC00282), showed an age-associated decrease in methylation in the PBMC EWAS and have not previously been reported to be associated with aging (Fig. 3 F,G, Supplementary table 4). Since PBMCs are made up of multiple cell types and the proportions can vary from sample to sample we used Meffil to estimate the proportions of cell types within each sample, and included the cell type proportions as covariates in a second EWAS calculation. FXR1 and ELOVL2 remained significant after cell type correction, however the association between cg10625583 (LINC00282) and age was no longer significant. The total number of significant sites identified was reduced, but it did not greatly affect the outcome of the PBMC EWAS (Supplementary Fig. 4). These results demonstrate that iPSCs do not retain age-related methylation signatures even outside of the sites used in epigenetic clocks, and EWAS may be used, even in a small sample size, to identify factors associated with aging in a cell type specific manner.

To provide an independent method for evaluating cellular age, we measured telomere length in all iPSCs. We found that telomere length was not correlated with donor age (Pearson r = - 0.0805, p-value = 0.4612) and appeared to be reset similarly to DNA methylation, consistent with previous reports (Supplementary Fig. 5) (Huang et al., 2011; Takasawa et al., 2018). Collectively, these data confirm that our iPSCs are epigenetically reprogrammed to a young state, in contrast to PBMCs from the same healthy donors across the human age span.

### methQTL in PBMCs and iPSCs are distinguished by cell state

The above data show that donor age does not contribute to the overall CpG methylation signal in iPSCs. Given that these cells are cultured under consistent conditions, any exposure-related methylation signals will likely be equivalent across lines, leaving genetics and cellular differentiation state as the most likely drivers of CpG methylation across the genome. We, therefore, evaluated patterns of genetically correlated CpG levels by identifying methylation quantitative trait loci (methQTL) in iPSCs and their donor PBMCs. Using tensorQTL (Taylor-Weiner et al., 2019), we considered *cis*-methQTL to include variants +/-1 Mb window to the CpG site, and identified 16,174 methQTL in PBMCs at a minor allele frequency (MAF) of >5% (Fig. 4A, Supplementary Table 5). As with the EWAS, we repeated methQTL analysis in PBMCs using cell type proportions as a covariate. Almost 72% of sites overlapped between both analyses (13,448) regardless of the covariate inclusion. Including cell type covariates resulted in 2615 new methQTL, and there were 2726 unique methQTL identified when celltype covariates were not used (Supplementary Table 6). Similarly, we recovered 8,013 methQTL in iPSC cells at >5% MAF (Fig. 4A, Supplementary Table 7). There were 1,092 methQTL that were statistically significant in both PBMCs and iPSCs. Examining these shared methQTL, we found that the effect sizes were generally consistent regarding direction and magnitude for both (Fig. 4B, Supplementary Table 8). However, a subset of CpG sites with methQTL detected in both PBMCs and iPSCs had opposite directions of effect (Fig. 4B). When comparing PBMC to iPSC, the majority of the significant methQTL (∼90%) were only found in one of the two assayed cell states.

**Figure 4.**
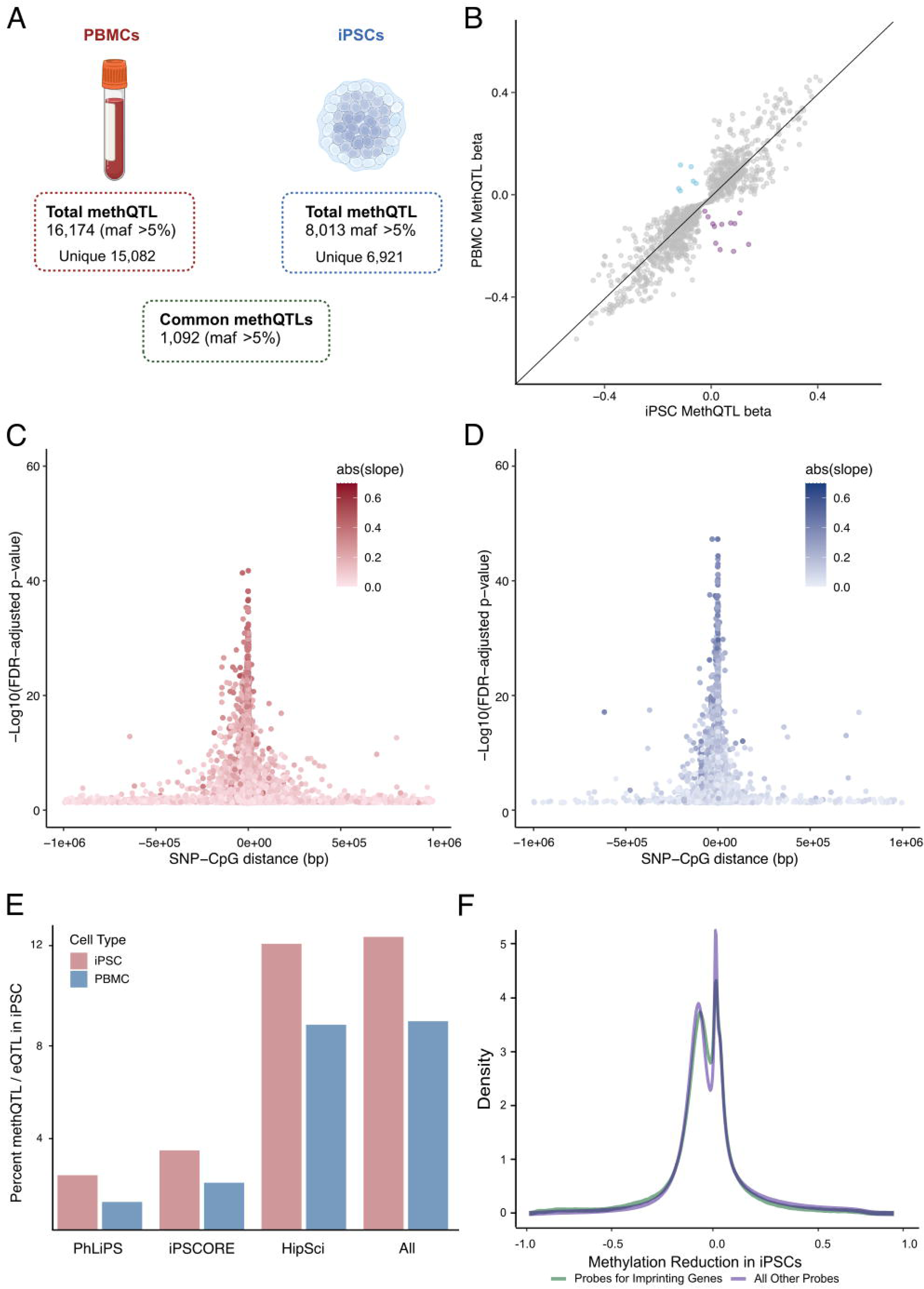
MethQTL analysis in PBMCs and iPSCs. **A**. Schematic representation of the workflow followed shows the number of methQTL that surpass the significant p-value threshold (FDR corrected p-value < 0.05) in PBMCs and iPSCs. Only CpG-SNP pairs with minor frequency allele MAF > 5% were retained for follow-up analysis. **B**. The scatter plot depicts the beta value of the significant methQTL in iPSCs (FDR corrected p-value < 0.05) on the x-axis and the significant methQTL in PBMCs (FDR corrected p-value < 0.05) on the y-axis. Blue dots represent CpG-SNP pairs, in which the alternative allele of the SNP increases the methylation of the CpG in PBMCs, and purple dots represent CpG-SNP pairs in which the alternative allele reduces the methylation of the CpG in PBMCs, while the effects are opposite in iPSCs. The distribution p-values of CpG-SNP pairs in PBMCs (**C, red**) and iPSCs (**D, blue)** with respect to the distance between features. Darker color indicates a stronger influence of genotype of methylation. **E.** Fraction of methQTL variants also identified as eQTL in published iPSC studies. **F.** Density of the change in methylation in PBMCs upon reprogramming into iPSCs separated by probes for imprinting genes (green) and all other probes (purple).

We saw more significant associations, i.e., methQTL with lower p-values, were localized to within 0.5 Mb of the CpG probe with which they were associated in both PBMCs and iPSCs (Fig. 4 C, D). Notably, very few methQTL probes were shared with CpG sites within the epigenetic clocks used to estimate epigenetic age (Supplementary Fig. 6A), further demonstrating that methylation at specific genomic sites is independently influenced by age or genetic sequence. There was a significant correlation between the beta estimates for methQTLs between PBMCs and iPSCs (Pearson correlation coefficient = 0.92), and the distribution of methQTL beta values was also similar between the different cell types (Supplementary Fig. 6 B,C). These results suggest that methQTL have similar overall properties between PBMCs and iPSCs, and in a number of cases the same SNP-probe pairs are congruent in both cell types.

Next, we hypothesized that methQTL may also act as eQTL due to the role of DNA methylation in the regulation of transcription. To answer this, we intersected methQTL from each cell type with eQTL previously identified in three large scale iPSC studies (Kilpinen et al., 2017; Panopoulos et al., 2017; Pashos et al., 2017). We found intersection of methQTL in both cell types identifying 1204 SNP-probe-gene sets in PBMCs and 850 SNP-probe-gene sets in iPSCs across all three datasets, however we show a significantly higher percentage of variants with QTL in both modalities in iPSCs (12.4% of iPSC methQTL) relative to PBMCs (8.8% of PBMC methQTL, χ =63.97, df=1, p < .00001) (Fig. 4E; Supplementary Table 9). Knowing that genetic imprinting occurs in the germline, we wondered if genetic imprinting also occurs during iPSC reprogramming. To test this, first we calculated the change in methylation at each probe from the differentiated PBMCs to the undifferentiated iPSC state for each donor. Then plotted the density of the change in methylation for the subset of probes that are known to be imprinted in humans relative to the change in methylation of all other probes (Fig. 4F). Based on this analysis there appears to be no enrichment in iPSC for methylation of probes associated with imprinting genes.

Querying annotated genes for methylation probes, methQTL signals that were significant only in PBMCs were enriched for genes that are involved in lipid localization and transport, and pathways associated with steroid secretion, while methQTL signals that were significant only in iPSCs are enriched for genes that are involved in microtubules and microtubule-associated processes (Fig. 5A). These analyses reinforce that despite methQTL having similar overall properties, the complement of methQTL that can be detected in each cell type are specific and related to cell function. To identify examples of the relationships that methQTL have between cell types, we examined the genotype:methylation relationship for probes near genes in either iPSCs or PBMCs. We identified a set of probes that shows consistent methQTL across both cell types with the same direction of effect in both cell types (Fig. 5B). The gene annotations for these probes (TMEM63B, PRR12, ERAP1, and two unannotated) identified genes that have low tissue specificity and are moderately expressed across many tissues. Next, we showed a set of methQTL SNP-probe pairs that are significant in PBMCs but not iPSCs (Fig. 5C) which include a SNP-probe pair associated with EGL9 Family hypoxia-inducible factor 1 (EGLN1). The cg18815120 site was fully methylated in iPSCs but showed genotype specific methylation in PBMCs. Interestingly, mutations in EGLN1 are linked to two different autosomal dominant blood disorders, familial erythrocytosis-3 and high altitude adaptation hemoglobin, reflecting an important role specific to blood cells (Lorenzo et al., 2014; Percy et al., 2006). We identified a final set of SNP-probe pairs that are significant in iPSCs but not PBMCs (Fig. 5D). The cg27577782 probe showed full methylation across genotypes in PBMCs but genotype specific changes in methylation in iPSCs. This site is associated with Ubiquitin-Specific Protease 36 (USP36) a gene that is 57% similar to a drosophila gene called scrawny that has been shown to be required in stem cell populations of multiple tissues for ubiquitin protease-regulated gene silencing (Buszczak et al., 2009). These examples suggest that the cell type specificity of some methQTL may contribute to complex phenotypes at the organismal level.

**Figure 5.**
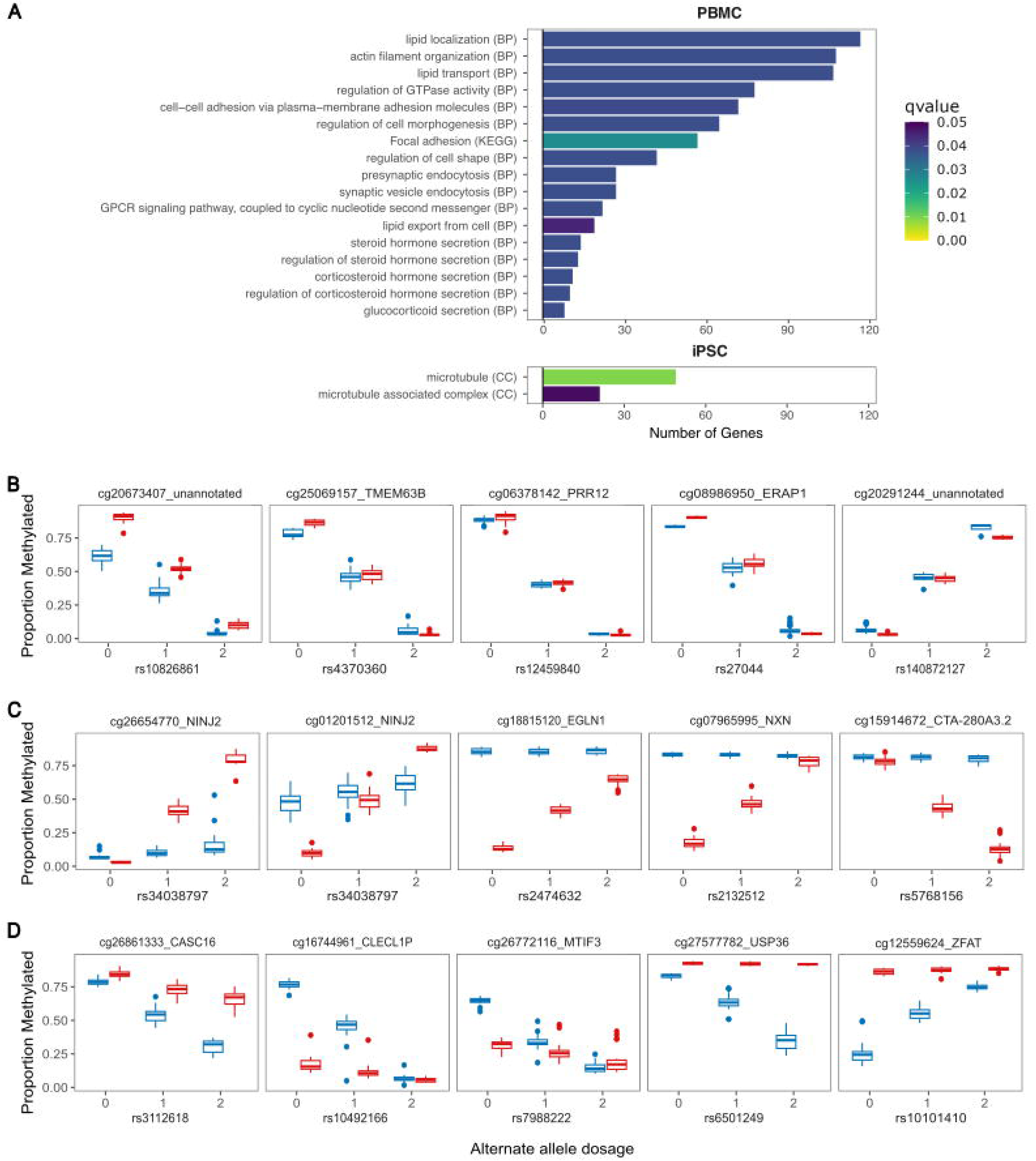
Cell type-specific methQTL enrichment. **A**. Gene Ontology terms identified for iPSC-specific methQTL (upper panel) and PBMC-specific methQTL (lower panel). Only biological process (BP) terms were found to be significantly associated with PBMCs, and only cellular component (CC) terms were associated with iPSCs. MethQTL were identified that show the same effect in both cell types (**B**), some that were PBMC-specific (**C,** red), and some that were iPSCs-specific (**D,** blue).

When cataloging enrichments, we noted that there was a greater proportional overlap in listed genes near methylation probes for which there was a methQTL than for methQTL overall. This suggested that, for any given locus, there might be multiple methQTL signals of which a proportion would be specific to a given cell type, extending the single examples discussed above. To evaluate this possibility, we identified loci for which there were more than ten methQTL with signals in either iPSCs and/or PBMCs that were annotated to the same unique gene. For all candidate loci (∼200), we plotted the mean methylation level in all samples for each cell type and annotated those which showed a significant methQTL signal in either iPSCs or PBMCs. We found that such loci have multiple methylation signals and that, as expected, a subset of those methylation signals were also methQTL, some of which were shared between cell types, but with a greater extent of cell-type specificity (Fig. 6). For example, at the *ADARB2*/*ADARB2-AS1* locus on chr10, which has been previously shown to be differentially methylated in Alzheimer’s disease (Konki et al., 2019), we identified methQTL-associated methylation in iPSCs that was fully unmethylated in PBMCs and vice-versa (Fig. 6A). We show additional examples near *B3GNTL1*, *HLA-DPB2*, and *SNTG2* (Fig. 6B-D). These results further support the concept that detection of a methQTL in a given cell type is facile when the underlying methylation feature shows intermediate methylation state in the cell population.

**Figure 6.**
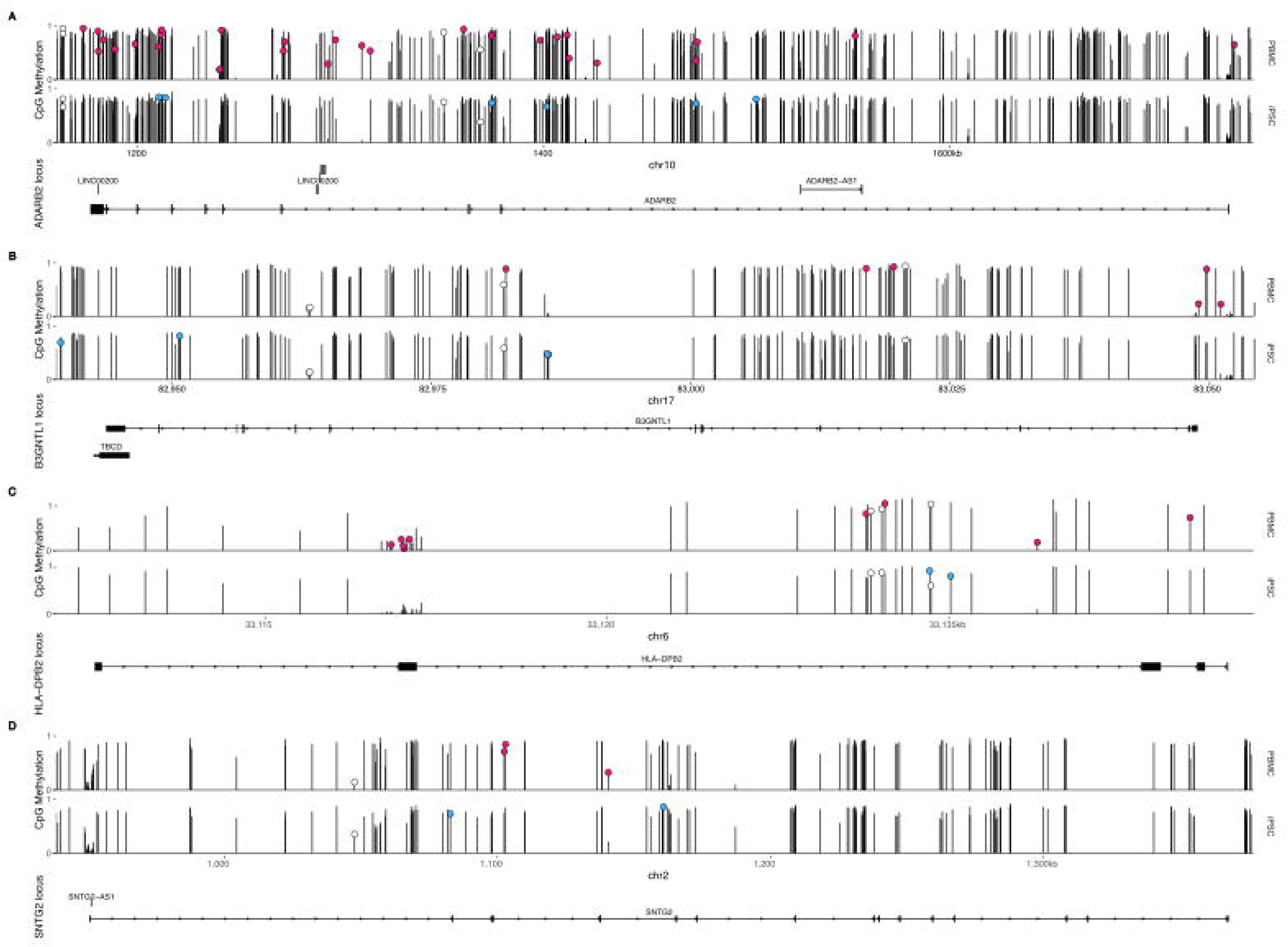
Genomic loci where multiple methylation signals can be detected but methQTLs show cell type specificity. Each plot shows methylation probes arrayed along the horizontal axis at the genomic locations on the indicated chromosome with average methylation values on the vertical axis for PBMCs (upper plots) and iPSCs (lower plots) samples. Methylation probes significantly associated with a neighboring genotype locus (methQTL) are labeled with circles, and are either PBMC-specific (red), iPSC-specific (blue), or present in both cell types (white). Examples are shown for four loci (**A**, *ADARB2*; **B**, *B3GNTL1*; **C**, *HLA-DPB2*; **D**, *SNTG2*) chosen from ∼200 candidates for informative density of signals over the selected genomic regions. Ideograms below each pair of methylation tracks show positions of exons for local genes for which the locus is named.

## Discussion

In this study, we set out to examine the effects of genetic variation in iPSCs relative to the PBMCs they were derived from on DNA methylation by measuring methQTL signals across the genome. We had the expectation that iPSCs are immature cells and hence lack age-related methylation changes, giving us the ability to isolate methQTL independent of aging. However, comparing PBMC methylation from the same donors, we find that methQTL are most strongly influenced by cell differentiation status.

To support these analyses, we generated a set of iPSCs from donors across human lifespan. Similar population-level collections have been reported in the past and have proven useful for genome-wide studies of human genetics and genomics at the endogenous level (Bonder et al., 2021; Bressan et al., 2023; HIPSCI Consortium et al., 2018). Our collection is characterized by the inclusion of donors without any clinical diagnoses at the time of collection and across a wide range of ages. We also included individuals with diverse ancestries; thus, this population partially reflects the community composition from which donors were recruited. Importantly for the current study, the same series of donors consented to genetic analysis, including genotyping and methylation from the PBMCs used to generate iPSCs, allowing for direct comparison between these two cell types.

Using DNA methylation, we confirm that iPSCs are epigenetically young compared to PBMCs from the same donors in regions where strong correlations between chronological age and methylation status were readily identified. This underlying result was stable across multiple clocks representing different sets of methyl-CpG sites across the genome (Hannum et al., 2013; Horvath, 2013; Levine et al., 2018; Zhang et al., 2019). Additionally, we used EWAS to evaluate age associations with methylation across the genome and noted that while there were many robust methylation signals in PBMCs, there were only a small number of nominally significant correlations in iPSCs. Several observations suggest that these candidate sites are likely not to be robust. First, given that we used a false discovery rate correction of 5%, the observed number of candidates (4) is lower than expected by chance sampling of ∼900,000 candidate sites. Second, estimated p-values for these four sites were only marginally above the significance threshold compared to the sites in PBMCs that greatly exceeded the same threshold. Third, examination of specific candidate CpG sites suggests that these are highly methylated in PBMCs and that any nominal correlation may be driven by a subset of iPSC lines with incomplete CpG methylation. Overall, these results demonstrate that iPSCs do not reflect the donor’s age from which they are derived, at least at the DNA epigenetic modification level.

Given that we have shown that iPSC generation removes the correlated effects of age on DNA methylation, and that the iPSC lines are maintained in a consistent environment, we therefore set out to examine genetic influences on CpG methylation by enumerating methQTL across the genome in both iPSC and PBMCs from the same donors. We find that methQTL share many characteristics between these two cell types, including an enrichment around gene bodies and proximity to the transcriptional start sites of genes similar to previous studies of methQTL in multiple tissues (Gibbs et al., 2010). Despite these general similarities in where methQTL are found across the genome, the actual sites that show methQTL are generally distinct between PBMCs and iPSCs. Specifically, while we identified ∼23,000 and ∼12,000 methQTL in PBMCs and iPSCs, respectively, only ∼3000 were shared between the two cell types, despite being derived from the same donors. Thus, while methQTL in different cell types may share similar generic properties, they are found in different genomic regions.

Examining which genes are associated with methQTL restricted to either cell type, we found that there were enrichments in PBMCs that reflect the function of these cells in the immune system. Furthermore, at many loci there were specific CpG methylation events that show genetic influence in only one cell type where intermediate methylation status was noted, i.e. that the methylation probe was neither fully methylated or fully unmethylated. Based on this data, we infer that DNA methylation is influenced by both cell type and genetic variation so that the two together affect measurable methQTL patterns. Specifically, we propose that for a methQTL to be measurable in a given differentiated cell type, the chromatin in that region must be permissive for transcription. A relationship between CpG methylation and changes in chromatin accessibility has been identified across species and tissues (Li et al., 2022), although it is not always clear whether methylation drives chromatin changes or vice versa. Because the current analysis focuses on genetic variation, we can reasonably conclude that methQTL are driven by genetic variants in each cell type but that the differences in methQTL between differentiation states are likely to be consequential to the establishment of cell identity, driven by cell type-specific transcriptional program and hence chromatin accessibility patterns. Future studies may take iPSCs and differentiate them into different target cell types, which could test this inference.

We acknowledge that there are several limitations to this study. First, while we could identify methQTL, our sample size is relatively small and would benefit from increased numbers to recover methQTL of smaller effect sizes. Second, we compared our iPSC data to PBMCs, which are heterogeneous cell populations and may underestimate the effects restricted to cellular subtypes. Additional confirmation in more restricted cell types would be helpful to exclude heterogeneity as a source of methQTL detection. However, this seems to be an unlikely confound, as we detected greater numbers of methQTL with biological enrichments in immune-related genes in the PBMCs than in the iPSCs that we would expect to be more homogenous in terms of cell type.

Overall, the current study attempted to disambiguate aging and cell type on CpG methylation state as defined by genetic variation captured in iPSCs. In contrast to our initial expectation, we find that methQTL are influenced strongly by cell type in that PBMCs could identify many more methQTL and many unique methQTL in functionally relevant gene enrichments compared to iPSCs. These results imply that methQTL detection may depend on cell type to a greater extent than has previously been appreciated.

### Experimental Procedures

#### Cohort Selection

PBMCs were collected from the Genetic and Epigenetic Signatures of Translational Aging Laboratory Testing Study (GESTALT) participants (n = 99) ages 22 – 92 years. As previously described (Roy et al., 2023, 2021; Tsitsipatis et al., 2023), GESTALT participants were clinically healthy at the time of enrollment based on eligibility criteria, they were not on medications except of one antihypertensive drug, free of major diseases except of history of silent cancer for more than 10 years. Participants had no physical or cognitive impairments, non-smokers and had a body mass index ≤ 30 kg/m2.

The GESTALT protocol was approved by the Institutional Review Board of the National Institutes of Health, and all participants provided written informed consent at each visit.

#### PBMC isolation

Peripheral blood mononuclear cells (PBMCs) were isolated from cytapheresis packs of the GESTALT donors by density gradient centrifugation using Ficoll-Paque Plus (Cytiva, Marlborough, MA). After subsequent washing steps with DPBS (Quality Biological, Gaithersburg, MD), PBMCs were cryopreserved as 10 million cells per ml and stored in a liquid nitrogen freezer until used.

#### PBMC DNA extraction and genotyping

DNA was extracted from 1 – 2 million PBMCs using either DNAQuik DNA extraction protocol (Reprocell, Beltsville, MD) or Qiagen DNeasy kit (Genetic Resources Core Facility, JHU, Baltimore, MD). 1 µg of DNA was sent to Novogene (Chula Vista, CA) for genotyping.

#### Reprogramming

PBMCs were sent to Thermo Scientific (Carlsbad, CA) for reprogramming. First, the cells were tested for mycoplasma, using the MycoSEQ™ kit (Applied Biosystems, 4460623). They were also confirmed to be free of human pathogens: hepatitis B virus (HBV), hepatitis C virus (HCV), Human Immunodeficiency Virus type 1 (HIV-1), Human Immunodeficiency Virus type 2 (HIV-2), Human T-lymphotropic virus type 1 (HTLV) I/II, Cytomegalovirus (CVM), Epstein–Barr (EBV), Herpes simplex virus 1 (HSV-1), and Herpes simplex virus 2 (HSV-2). If any lines tested positive for any of these, they were excluded from reprogramming. Mycoplasma and human pathogen-negative PBMCs were thawed, assessed for viability, and transduced with CytoTune^TM^ reprogramming particles from the Cytotune^TM^-iPS 2.0 Sendai Reprogramming kit (Invitrogen, A16518). Three days after transduction, the cells were re-plated onto Geltrex™ (LDEV-Free hESC-qualified Reduced Growth Factor Basement Membrane Matrix)-coated dishes (Gibco, A1413202). The cells were maintained in Essential 8™ medium (Gibco, A1517001), which was changed daily until colonies were picked for further analysis.

#### Colony selection

Four to five weeks after introducing Cytotune^TM^ 2.0 particles, pluripotent stem cell-like colonies emerged and were picked based on morphological characteristics. Colonies were cut out using a needle and plated into a well of Geltrex™-coated 24-well plates containing Essential 8™ Medium. Colonies were expanded, and then dissociated using TrypLE™ Select Enzyme (Gibco, 12605010) and single cell passaged into one well of a Geltrex™-coated 12-well plate containing StemFlex™ Medium (Gibco, A3349401), supplemented with RevitaCell™. Once established in 12-well plates, the two best-looking colonies were dissociated using Versene Solution (Gibco, 15040066) and plated in Geltrex™-coated 6-well plates. The cell lines were then expanded and banked for liquid nitrogen storage using the PSC Cryopreservation kit (Gibco, A2644601). Upon expansion, cell pellets were collected for Mycoplasma testing, TaqMan® iPSC Scorecard™ (ThermoFisher, A15872), Pluritest™ (Thermo Scientific, A38154), and Karyostat+™ (Thermo Scientific, A52986).

#### Culture of iPSCs

All resulting iPSC lines were expanded in Essential 8™ Flex medium (Gibco, A2858501) on Matrigel (Corning, 354230)-coated dishes. RevitaCell (Gibco, A2644501) supplement was added to the medium for one day after thawing and cell splitting. iPSCs were passaged using TrypLE™ Select Enzyme or Cell Dissociation Buffer (Gibco, 13151014) for a total of 1-2 passages and recovered post-split using Essential 8™ Flex Medium with 1X RevitaCell™. Expanded iPSCs were frozen for storage using Synth-a-Freeze™ Cryopreservation Medium (Gibco, A1254201).

### DNA isolation and genotyping iPSCs

Approximately 1×10^6^ cells were pelleted and frozen for DNA isolation. Genomic DNA was extracted from iPSCs using the Maxwell RSC Tissue DNA kit (Promega, AS1610) following the manufacturer’s instructions. DNA samples were quantified, and 250 ng of genomic DNA were genotyped for each sample using the Neuro Global Diversity Array-8 v.1 LCG with Neuro Booster content (Illumina, 20031815 and 20042459) according to the manufacturer’s instructions. Briefly, genomic DNA was amplified and enzymatically fragmented. Following alcohol precipitation, the resulting fragments were resuspended and hybridized to the array.

Automated allele-specific, enzymatic base extension and fluorophore staining were performed using a liquid-handling robot (Tecan, Research Triangle Park, NC). The stained arrays were washed, sealed, and vacuum-dried before scanning on the Illumina iScan instrument.

### Genotyping quality control and imputation for iPSCs

Raw data files were imported into the GenomeStudio Genotyping Module (v.2.0, Illumina) using a custom-generated sample sheet, and genotypes were called using a GenCall threshold of 0.15. The genotype data were exported in a ped-file format for downstream quality control. Genotype quality control was performed following standard criteria. Samples meeting the following criteria were excluded: (1) samples with inbreeding coefficient F-statistic (estimated by PLINK) outside of the range (−0.15, 0.15); (2) low call rate (≤ 95%); (3) mismatch between reported sex and genotypic sex; (4) duplicate samples (pi-hat statistics > 0.8). Ancestry was ascertained based on principal component analysis compared to the HapMap 3 Genome Reference Panel. No samples based on ancestry were excluded from the follow-up analysis. Related samples were also included for imputation (defined as having a pi-hat > 0.125).

Variants-wise quality was also applied, and variants were removed as follows: (1) monomorphic SNPs; (2) palindromic SNPs; (3) variants with haplotype-based nonrandom missingness (*p* ≤ 1.0 × 10^−4^); (5) variants with an overall missingness rate of ≥ 5.0%; (6) non-autosomal variants; and (7) variants that deviated from Hardy-Weinberg equilibrium (*p* ≤ 1.0 × 10^−10^). Imputation was performed against the Trans-Omics for Precision Medicine (TopMed) imputation reference panel for hg38 via minimac4 using data phased by Eagle v2.4 provided by TopMed Imputation Server (33). After imputation, only variants with an imputation quality score (R^2^) > 0.8 and minor allele frequency > 0.01 were included in the analysis.

### DNA methylation profiling

For each sample, 450 ng of genomic DNA was bisulfite converted (Zymo Research, D5003), hybridized onto Infinium HumanMethylation EPIC BeadChips v1.0 (Illumina, 2016168), and stained according to the manufacturer’s protocol. Briefly, after DNA amplification at 37°C for 20 hours, the samples were enzymatically fragmented and precipitated, followed by DNA hybridization and automated Beadchip staining using a liquid handling robot (Tecan).

Methylation arrays were washed, sealed, and vacuum-dried before scanning using the iScan instrument (Illumina).

### Methylation quality control and normalization

Methylation data quality control and normalization was performed using the meffil R package (version 1.3.3) (Min et al., 2018). Samples meeting the following criteria were excluded: (1) samples with a predicted median methylated signal > 3 standard deviations (SD) from the expected, (2) samples that had > 10% of probes with bead numbers < 3, (3) samples that had > 10% of probes with detection *p*-value > 0.01, (4) samples with gender mismatch between the reported and predicted gender (sex outlier value > 5 SD from the mean), (6) samples with a genotype concordance < 80% (genotype mismatch). Probe-wise, we excluded CpGs within single nucleotide polymorphisms with a minor allele frequency of >1% located within five nucleotides of the target sites, probes tagging non-unique 3΄ bases long, and cross-reactive probes as previously described (Chen et al., 2013). Additional EPIC-specific cross-reactive probes were obtained from the Maxprobes R package (version 0.0.2) and removed (McCartney et al., 2016; Pidsley et al., 2016). Principal component analysis was performed on the control matrix and the 20,000 most variable probes.

### Epigenetic clock analysis

The calculation of the epigenetic age using the Horvath (Horvath, 2013), Hannum’s (Hannum et al., 2013), phenoAge (Levine et al., 2018), BLUP (Best Linear Unbiased Prediction) (Zhang et al., 2019), and EN (Elastic Net) clocks were performed with the Methylclock R package (version 1.6.0)(Pelegí-Sisó et al., 2021).

### Analysis of absolute telomere length

The average telomere length for each reprogrammed iPSC line was determined using the Absolute Human Telomere Length Quantification qPCR Assay Kit (Sciencell, 8918). Briefly, 5 ng of DNA was used for each reaction with Single Copy Reference (SCR) primers or Telomere (TEL) primers. Each iPSC line was measured in technical triplicate for both primer pairs. A reference human genomic DNA sample of known length (935 ± 45 kb per diploid cell; Sciencell, 8918b, Lot #33114) was included in triplicate with both primer pairs. The relative telomere length and total telomere length of each iPSC line was determined as the protocol directed using the ^ΔΔ^Cq method (Livak and Schmittgen, 2001). The average telomere length on each chromosome end (telomere length per diploid cell divided by 92 chromosome ends in one diploid cell) was plotted against the age of the donor when PBMCs were isolated for reprogramming. Correlation between donor age and average telomere length identified the Pearson r = −0.0805 (p-value = 0.4612).

### PBMC cell type predictions

To determine if cell type heterogeneity within PBMC samples impacted EWAS and methQTL, we performed these analyses with and without covariates of cell type proportions. To generate covariates, we used the R package Meffil’s meffil.estimate.cell.counts.from.betas function (Min et al., 2018). Using data from (Bakulski et al., 2016) as a model, the function predicts cell type proportion for seven cell types (B, CD4T, CD8T, granulocytes, monocytes, natural killer lymphocytes, and nucleated red blood cells) based on methylation beta values for each sample.

### Epigenome-wide association study

Epigenetic genome-wide association analyses were performed independently in PBMCs and iPSCs using the *meffil.ewas* function of the R package meffil (v 1.3.3) (Min et al., 2018). Linear regression was performed at each site (n = 748,760 probes in the PBMC dataset and n = 732,148 probes in the iPSC dataset) testing for an association between DNA methylation β values and the donor’s age at the time of the sample collection (age at collection). For covariates, the model included the sex, first five principal components from LD-pruned genotypes (Manichaikul et al., 2010), and the first five principal components from the top 20,000 most variable methylation probes. For PBMCs, analysis was also repeated with cell type proportion predictions as covariates.

### methQTL analysis

Genome-wide SNP-methylation probe pairs were tested using TensorQTL, a GPU-enabled QTL mapper that achieves >200 fold faster *cis-*QTL calculation than the CPU-based mappers (Taylor-Weiner et al., 2019). Briefly, an additive linear model tested if genotypes within 1MB of a probe site (*i.e.* the number of alternative alleles) predicted DNA methylation. iPSC and PBMC methylation datasets were processed independently and included for covariates sex, age, and the first ten principal components from LD-pruned genotypes. For PBMCs, analysis was also repeated with cell type proportion predictions as covariates.

### Gene Ontology Enrichment Analysis

MethQTL from the previous analysis were filtered to datasets that were unique to each cell type by removing any overlapping sites. Genes from each set were run using gprofiler2_0.2.2 within R with default parameters including using the g:SCS algorithm to adjust enrichment p values for multiple testing (Raudvere et al., 2019).

## Supporting Information

Methylation beta predictions for samples and probes (following quality control) is available at Zenodo (https://doi.org/10.5281/zenodo.15191371). Code associated with this analysis is available on GitHub (https://github.com/neurogenetics/methylation-iPSC-PBMC).

## Acknowledgements

This research was partly supported by the Intramural Research Program of the NIH, the National Institute on Aging (ZIAAG000931-03 to MRC), and the National Institute of Neurological Disorders and Stroke (ZIANS003154 to SWS). This work utilized the computational resources of the NIH HPC Biowulf cluster (http://hpc.nih.gov). GESTALT study participants were recruited under National Institute on Aging IRB protocol 15-AG-0063. Written consent was obtained from each individual.

