## Supplementary Figures for "Characterization of DNA methylation in PBMCs and donor-matched iPSCs shows methylation is reset during stem cell reprogramming"

**Contents**

Supplementary Fig. 1: iPSC lines are generated from a cohort of healthy donors.

Supplementary Fig. 2: Correlation between clones and between donors.

Supplementary Fig. 3: Correlation matrix between PBMCs and iPSC clones.

Supplementary Fig. 4: EWAS in PBMCs adjusted for cell type proportions.

Supplementary Fig. 5: Telomere length in iPSC.

Supplementary Fig. 6: Overlap of epigenetic aging clock and methQTL CpG methylation sites.

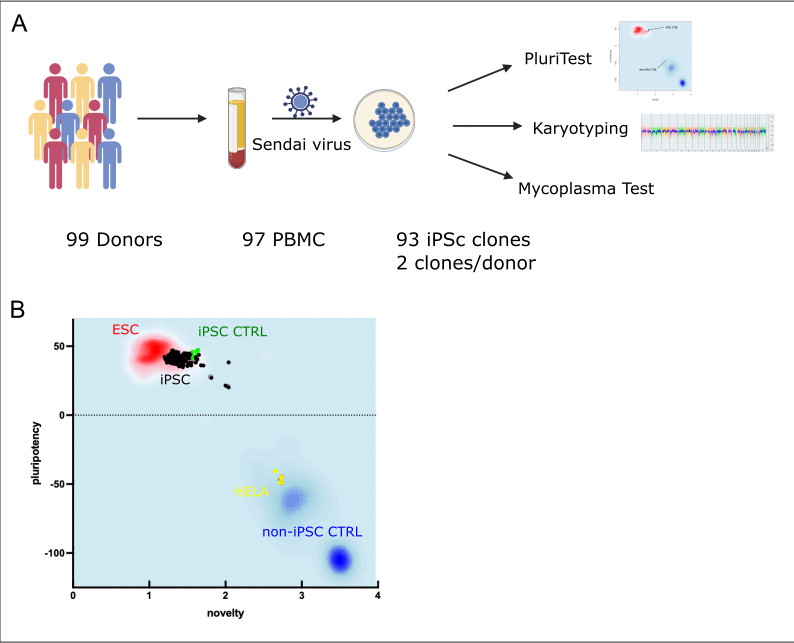

**Supplementary Figure 1.** iPSC lines are generated from a cohort of healthy donors. After blood collection from a healthy cohort of donors, PBMCs are reprogrammed into iPSCs following quality control measures. **A**. Schematic diagram of the workflow used for iPSCs generation. PBMC samples from 99 donors were reprogrammed using CytoTune-iPS 2.0 Sendai Reprogramming Kit. Two clones per donor were generated, and a PluriTest characterization was applied to each clone. The PluriTest characterization assay includes 1) karyotyping, using the array-based Applied Biosystems™ KaryoStat™; 2) pluripotency, using both the Pluritest that consisted of the analysis of 36,000 transcripts and variants to test pluripotency and the Scorecard, that consisted on a TaqMan array of 93 genes to confirm pluripotency and differentiation potential; and 3) mycoplasma testing, using the MycoSEQ Mycoplasma qPCR Detection Kit. **B**. The Pluritest results are represented in a pluripotency plot. The x-axis shows the novelty score results, while the y-axis represents the pluripotency score results per each clone. The red and blue background depicts the empirical distribution of the pluripotent (red) and non-pluripotent (blue) samples in the reference data set of 223 hESCs, 41 iPSCs, and 186 somatic cell tissues. The distribution of the tested iPSc samples can be seen in black. A non-iPSC sample derived from HELA cells ( yellow) was included in this experiment to serve as a negative control for non-pluripotency.

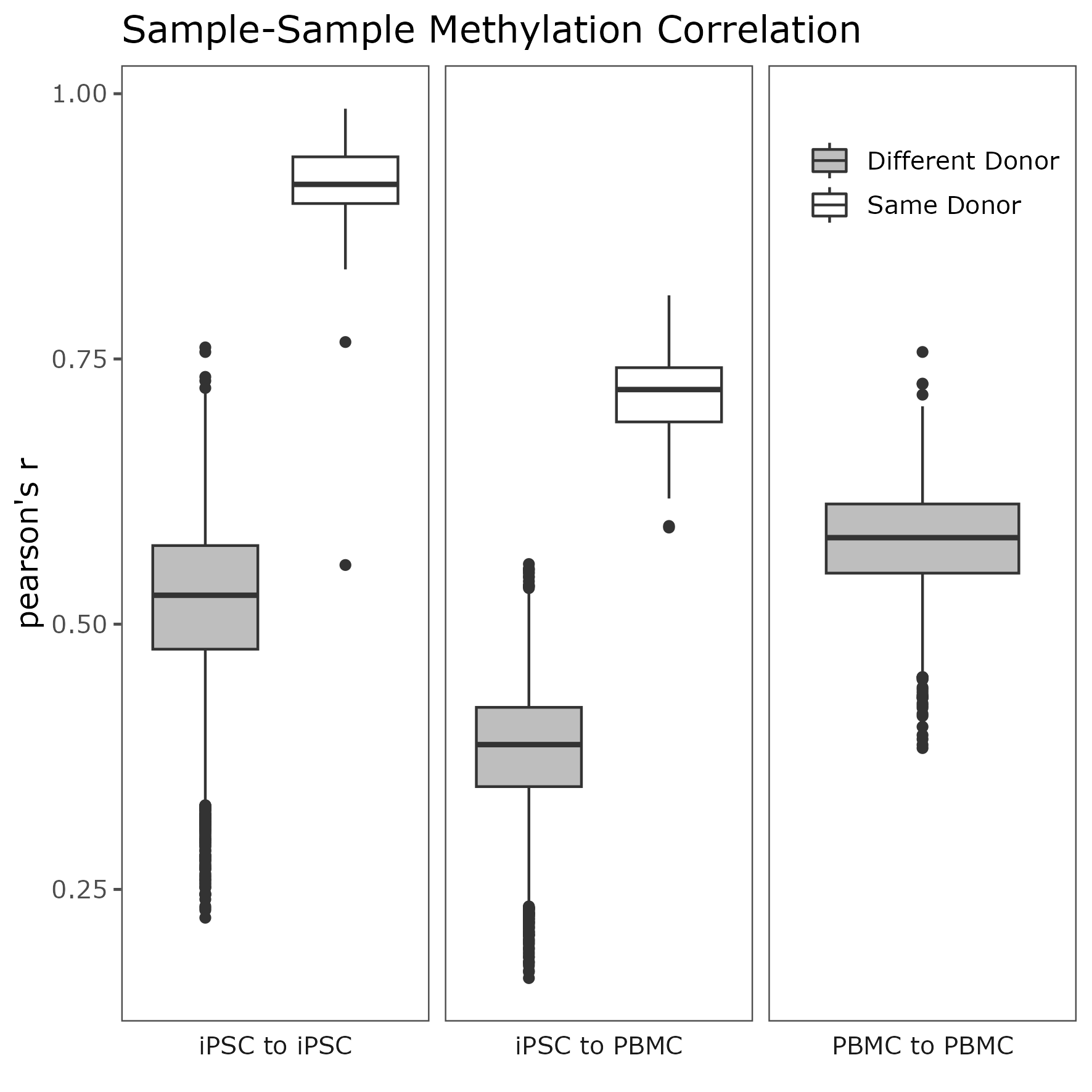

**Supplementary Figure 2. Methylation variability between clone-to-clone and PBMC-to-clone comparison.** The box plot depicts the result of the Pearson correlation coefficient comparison between iPSC clones from the same donor (white), same donor PBMC to iPSC clones (gray), and unrelated donors (including both clone to clone and PBMC to clone comparison). The group comparison is on the x-axis, while the y-axis represents the Pearson correlation coefficients per group. All group comparisons were significantly different from each other (ANOVA: *df*=4, *F*=10320, *p*<2×10^-16^; Tukey HSD, all comparisons: *p*<2×10^-16^).

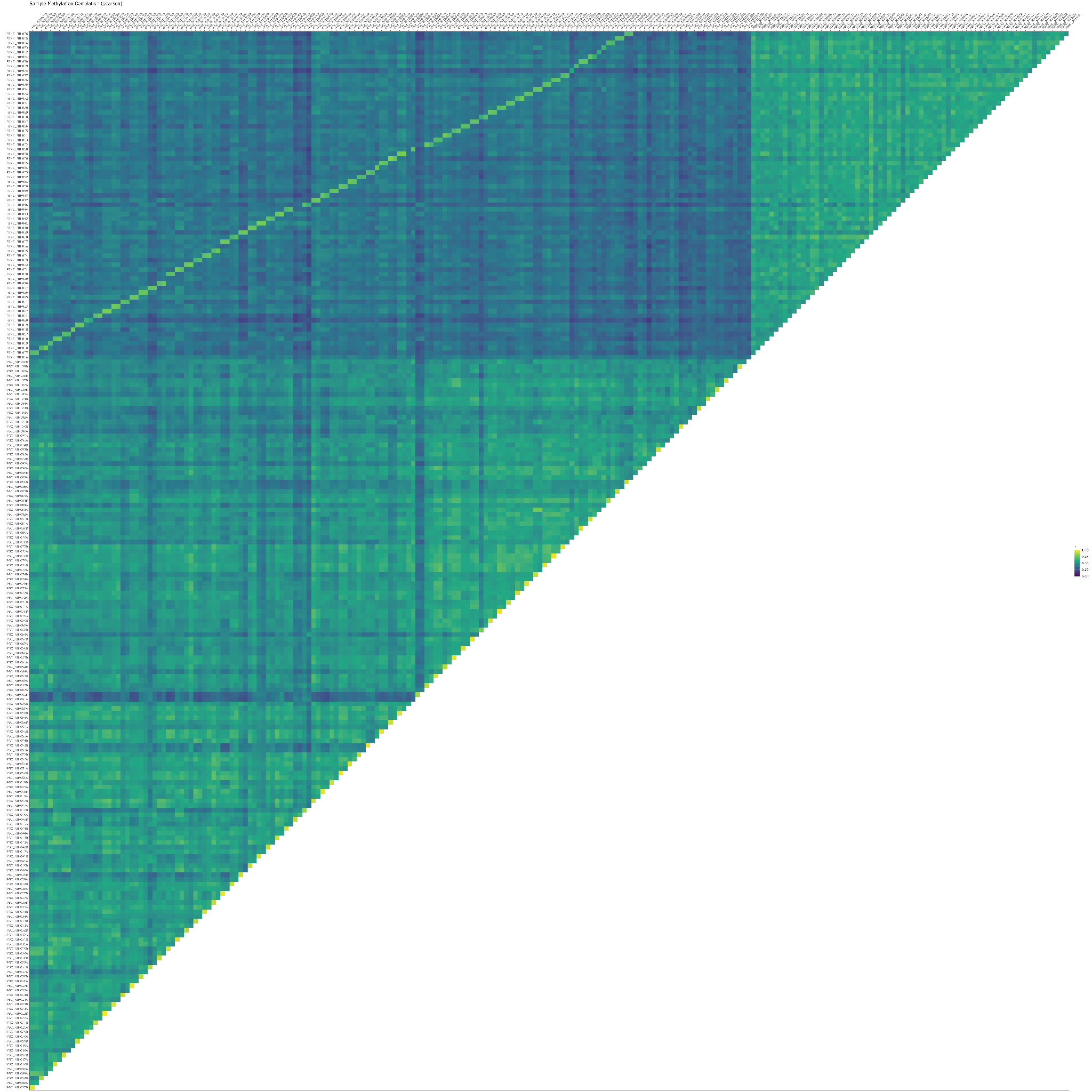

**Supplementary Figure 3. Correlation matrix between PBMCs and iPSC clones.** Pearson correlation analysis was performed using 775 informative methylation probes. Informative methylation probes were selected by intersecting the top 5000 most variable probes for iPSCs and PBMCs.

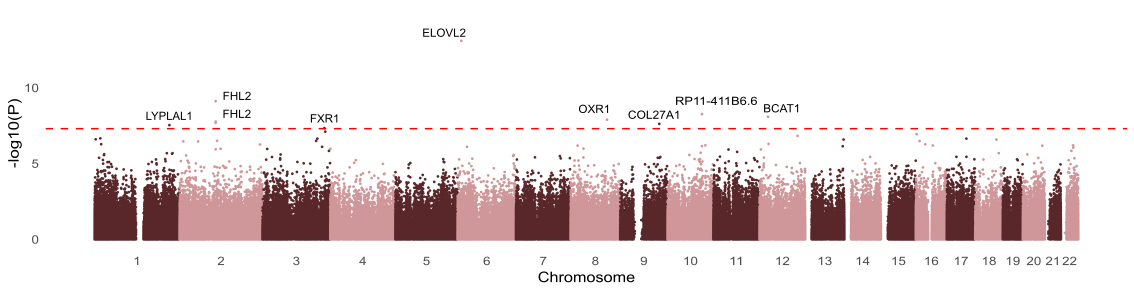

**Supplementary Figure 4. EWAS in PBMCs adjusted for cell type proportions.** Manhattan plot showing significant associations between DNA methylation at common CpG sites and age in cell type proportion adjusted PBMCs.

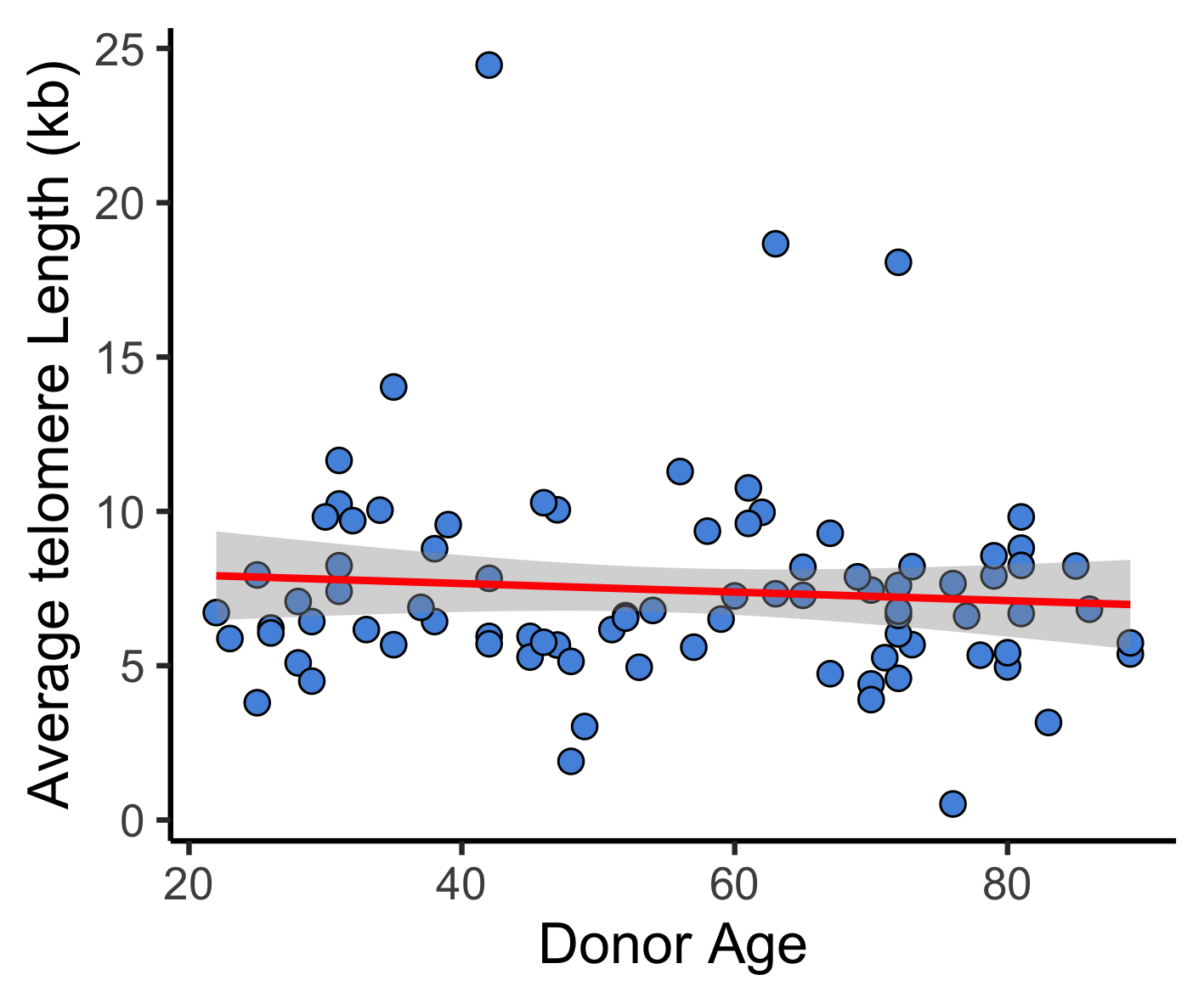

**Supplementary Figure 5. Telomere length in iPSCs.** Correlation between donor age at the time of collection (x-axis) and [Telomere Restriction Fragment](https://www.ncbi.nlm.nih.gov/pmc/articles/PMC4972328/) TRF length (y-axis) in iPSC. Telomere length is not correlated to biological age at collection in iPSCs, r^2^=−0.0065.

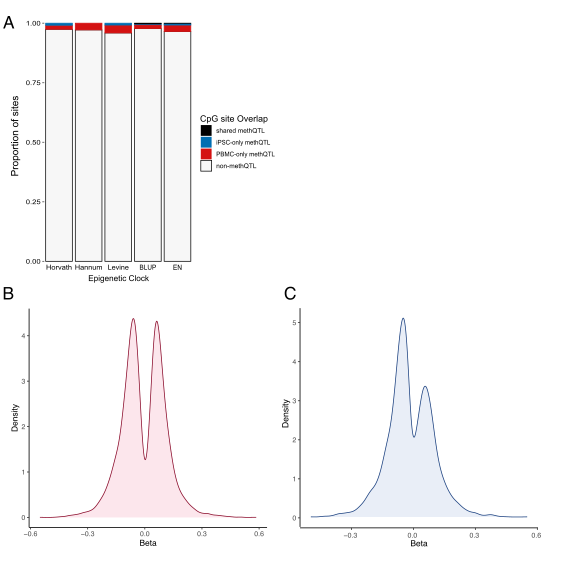

**Supplementary Figure 6. Overlap of sites in methylation clocks with methQTL**. For each of the clocks listed on the horizontal axis, we plotted the proportion of methylation sites that were identified as significant methQTLs in iPSCs (blue), PBMCs (red), both cell types (black), or not significant in either cell type (gray). The majority of sites were not significant methQTL sites for either cell type. **B,C**. The plots show the beta distribution of the significant CpG-SNP pairs identified in PBMCs (red, **B**) and iPSCs (blue, **C**).
